# Sex-specific dominance maintains multilocus polymorphism and can resolve sexual conflict

**DOI:** 10.64898/2026.09.09.750493

**Authors:** Mattias Siljestam, Göran Arnqvist, Claus Rueffler

## Abstract

Sexually antagonistic selection arises when females and males favour different phenotypes but share their genetic basis. Beneficial sex-specific dominance can alleviate this conflict by allowing each allele to be more dominant in the sex in which it is beneficial. Whether the coevolution of dominance and allelic diversity can generate and maintain polymorphism across a polygenic trait remains unclear. We investigate a quantitative trait encoded by multiple diploid loci, with allelic effects shared between the sexes and evolving sex-specific promoter affinities determining dominance. We show that sex-specific dominance can generate polymorphism across many loci, including under weak or asymmetric selection. Whereas additive allelic effects lead to the concentration of polymorphism at a single major locus, evolving dominance allows polymorphism to become distributed across loci. This distributed polygenic architecture generates sexual dimorphism while reducing within-sex phenotypic variance: increasing the number of polymorphic loci, as well as allelic diversity within loci, allows female and male phenotypes to approach their respective optima with lower segregation load. Sufficiently strong selection can instead cause polymorphism to collapse toward a single major locus, but the selection strength required for this transition increases sharply with the number of contributing loci. Thus, evolving sex-specific dominance can make widespread genetic polymorphism part of the resolution of sexual conflict.

## Introduction

Sexually antagonistic (SA) selection arises when the optimal phenotype differs between females and males. Because the two sexes share much of their genetic basis, this can generate genetic conflict in which different alleles are favoured in the two sexes. A large body of theory has elucidated under what conditions SA selection favours genetic polymorphism, maintaining several segregating alleles within a population (reviewed in Flintham et al., 2026). Single-locus models show that SA selection can maintain two alleles, one favoured in females and one in males, when selection in the two sexes is sufficiently strong and symmetric (e.g., Kidwell et al., 1977; Flintham et al., 2024; Siljestam et al., 2024). The conditions for coexistence are substantially relaxed when dominance is sex-specific, such that each allele contributes more strongly to the phenotype in the sex in which it is more beneficial (Kidwell et al., 1977; Flintham, 2025; Siljestam et al., 2024). In this case, polymorphism can also be maintained under weak or asymmetric selection. Whether such dominance itself can readily evolve is therefore central to understanding the maintenance of sexually antagonistic variation.

Classical theory emphasizes that dominance evolves particularly readily when balancing selection maintains polymorphism, because dominance modifiers affect fitness through heterozygotes (for a review see Otto and Bourguet, 1999). Consistent with this view, Spencer and Priest (2016) showed theoretically that sex-specific dominance evolves if a polymorphism maintained by SA selection exists in the first place. In a recent modelling study, however, we showed (Siljestam et al., 2024) that the conditions for the evolution of sex-specific dominance, and thereby for the evolution and maintenance of genetic variation at a locus under SA selection, are much broader. Our model considers a quantitative trait encoded by a single diploid locus under the continuum-of-alleles model (Kimura, 1965), where each allele is associated with a promoter affinity determining its dominance in the two sexes relative to other alleles. Coevolution between allelic values and these sex-specific promoter affinities can generate a positive feedback in which beneficial sex-specific dominance turns stabilizing selection on allelic values into disruptive selection. Divergence in allelic values, in turn, strengthens selection for further sex-specific dominance. This allows heterozygote phenotypes to approach the sex-specific optima, reducing segregation load. Importantly, this feedback does not require a balanced polymorphism to be present initially. It can begin during selective sweeps, during evolutionarily transient protected polymorphisms, or from variation generated by mutation– selection–drift balance (Siljestam et al., 2024), extending the classical emphasis on dominance evolution within an already maintained polymorphism. Moreover, more than two alleles can evolve and coexist, with sex-specific dominance hierarchies emerging within such polyallelic polymorphisms.

Single-locus models are convenient but may provide a poor approximation for quantitative traits encoded by many loci. It is therefore natural to ask whether these single-locus dynamics persist under a polygenic architecture. Three existing studies investigate the evolution of polygenic traits under sexually antagonistic selection, but address somewhat different questions. First, Flintham et al. (2024) consider a multilocus model with two possible alleles per locus and with alleles acting additively within and across loci. Under these assumptions, the authors find that, similarly to the single-locus case, considerable genetic variation is maintained only under strong and symmetric selection.

Second, Flintham (2025) extend this model by allowing evolving modifiers to alter either sex-specific dominance (as in Siljestam et al. 2024) or sex-specific expression. In the first case, they find that a multilocus architecture significantly slows down the evolution of dominance modification as selection becomes weaker. In the second case, sex-specific expression provides a route to architectures in which the two sexes use different subsets of loci. Such an architecture ultimately allows a genotype homozygous at every locus to produce the optimal female phenotype in females and the optimal male phenotype in males, and thus completely resolves the conflict arising from sexually antagonistic selection. However, this result hinges on two strong assumptions. First, even at the lowest redundancy considered, the sex-specific optimal phenotypes can be produced by only half of the loci. Second, loci can be silenced in one sex without disrupting other essential functions that require expression of the same genes in that sex.

The third study, Flintham et al. (2026), considers a polygenic architecture in which evolution at each locus follows a continuum-of-alleles model (Kimura, 1965). Two types of mutations are allowed: mutations in dominance modifiers alter sex-specific dominance, whereas mutations affecting the trait allele directly alter its phenotypic effects in females and males. Although these mutational effects are generally positively correlated between the sexes, repeated mutation allows the same allele to accumulate different phenotypic effects in females and males. Selection can therefore produce combinations of alleles whose summed effects approach the female optimum in females and the male optimum in males. As in the sex-specific-expression model of Flintham (2025), a genotype homozygous at every locus can therefore resolve the genetic conflict.

Thus, the latter two studies both allow the sexual conflict to be resolved without maintaining allelic polymorphism: in one case through sex-specific expression of loci, and in the other through sex-specific effects of the alleles themselves. If these routes evolve to completion, the conflict that could maintain sexually antagonistic polymorphism is removed, even when sex-specific dominance can evolve.

A parallel body of theory has investigated the evolution of polygenic quantitative traits under diversifying selection generated by ecological heterogeneity (Kopp and Hermisson, 2006; van Doorn and Dieckmann, 2006). These studies predict that genetic variation maintained by balancing selection and initially distributed over many loci becomes concentrated at a single major locus when selection favours two distinct phenotypic optima. Under additive allelic effects, such concentration reduces the production of maladapted intermediate phenotypes and thus reduces segregation load. This raises the question of whether such a concentration of allelic effects could also occur for a polygenic trait under sexually antagonistic selection when sex-specific dominance is allowed to evolve.

In this study, we generalize the model of Siljestam et al. (2024) from one locus to a polygenic architecture. We allow for up to 30 independently segregating loci to contribute additively to a quantitative trait under sexually antagonistic selection, while alleles within each locus can evolve sex-specific dominance through changes in their promoter affinities. Unlike Flintham (2025), allelic evolution at each locus follows a continuum-of-alleles model and is therefore not restricted to two alleles. In our model, we explicitly exclude the possibility of sex-specific or sex-limited allelic effects that lead to the erosion of genetic variation in the studies by Flintham (2025) and Flintham et al. (2026). There are two main reasons for this exclusion. First, many genes are pleiotropic and perform essential functions, constraining the evolution of sex-specific allelic effects. Consequently, sex-limited expression or strongly sex-specific effects may be deleterious because the same gene product is required for essential functions in both sexes (Mank and Ellegren, 2009). Second, empirical data show that genetic correlations between homologous traits expressed in males and females are generally large and positive, demonstrating that allelic effects are in fact typically shared between the sexes (Poissant et al., 2010).

Under this constraint, we find that beneficial sex-specific dominance qualitatively changes the evolution of the polygenic architecture. Rather than concentrating polymorphism into a single locus (as predicted by Kopp and Hermisson, 2006; van Doorn and Dieckmann, 2006), SA selection generates beneficial sex-specific dominance and polymorphism across all loci, often consisting of more than two alleles per locus. Increasing both the number of loci across which phenotypic effects are distributed and the number of alleles maintained within loci results in increasingly pronounced sexual dimorphism, with phenotypic distributions becoming progressively narrower around their respective optima. The sexual conflict can therefore be at least partly resolved while extensive genetic variation is maintained.

### The model

We consider a panmictic population of constant size *N* with non-overlapping generations Fisher (1930); Wright (1931). Each offspring is produced by independently sampling one female and one male parent with probabilities proportional to their reproductive success. Each parent contributes a gamete through fair meiosis and free recombination among loci, and the two gametes fuse to form a zygote. This process is repeated until *N* offspring constituting the next generation have been produced.

We investigate the evolution of a quantitative trait, denoted *z*, that is under sexually antagonistic (SA) selection. We assume that *z* determines fitness through reproductive success, and it could represent any shared characteristic, such as morphology, colouration, physiology, behaviour, or a life-history trait. Female reproductive success can, for example, represent the number of eggs or offspring produced, whereas male reproductive success can represent mating and fertilization success.

The reproductive success (fitness) of an individual of sex *k* ∈ {♀,♂} with phenotype *z*_*k*_ is given by the Gaussian function

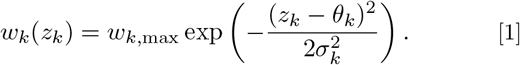

Here, *θ*_*k*_ is the sex-specific phenotypic optimum and *σ*_*k*_ determines the width of the fitness function. Reproductive success matters only relative to that of other individuals of the same sex, so *w*_*k*,max_ cancels and therefore can be set to one without loss of generality.

We place the female and male optima at *θ*_♀_ = *δ/*2 and *θ*_♂_ = *δ/*2, such that *δ* gives the distance between the sex-specific optima. The widths of the Gaussian fitness functions, *σ* and *σ* determine the strength of selection in each sex, with larger values of 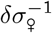 and 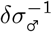 corresponding to stronger selection relative to the distance between the optima. For presentation purposes, we set *δ* = 1 in the simulations and therefore report selection strength using 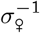 and 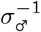 throughout the main text. Figure 1 illustrates how increasing selection strength changes the trade-off between female and male fitness; the weak and strong cases, *σ*^−1^ = 1 and *σ*^−1^ = 3, correspond to the selection strengths used in our focal simulations below.

**Figure 1.**
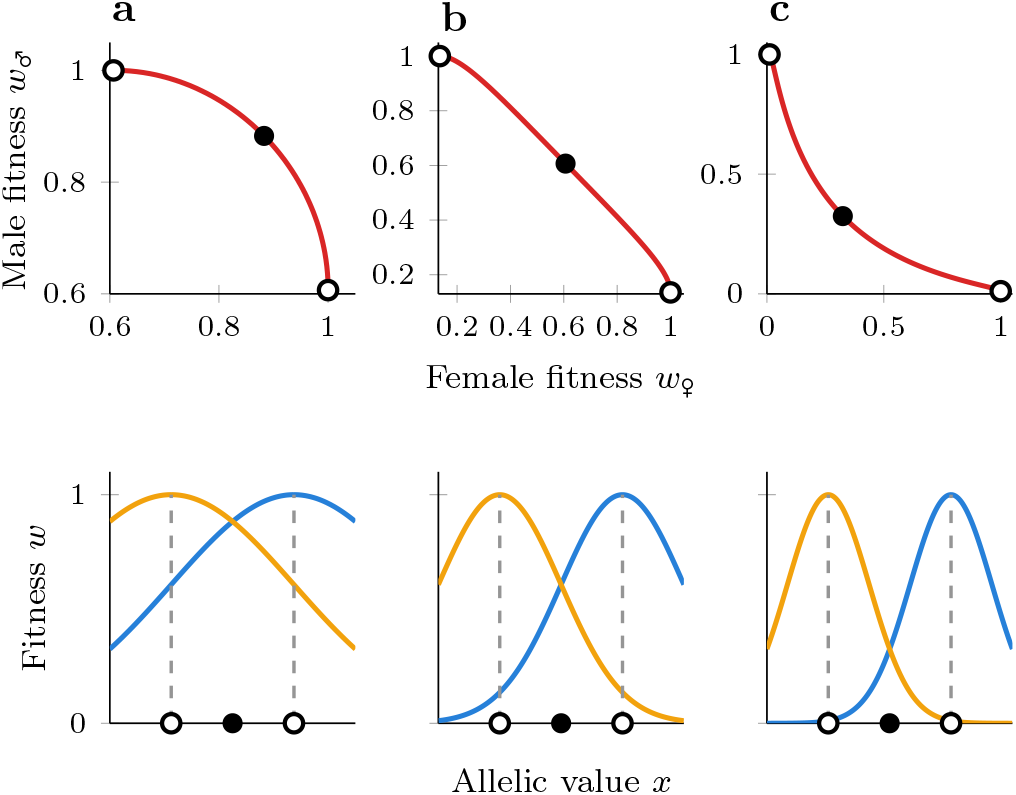
Trade-off between female and male fitness (reproductive success). The upper panel illustrates the trade-off as a parametric curve of the underlying phenotypic trait value *z*, while the lower panel shows the underlying Gaussian fitness functions for females and males in blue and orange. The sex-specific phenotypic optima are indicated by open dots, and the singular phenotype *z*^*∗*^ by the filled dot. In **a**–**c**, Gaussian functions with decreasing widths (*σ*^*−*1^ = 1, 2 and 3) are illustrated, resulting in a weak, locally linear (around *z*^*∗*^), and strong trade-off, respectively.

The trait is encoded by *L* independently segregating autosomal loci. The phenotype expressed by an individual of sex *k* is determined by the additive contributions from all loci,

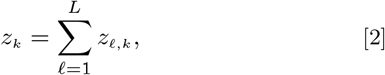

where *z*_*ℓ,k*_ is the contribution of locus *ℓ* to the phenotype expressed in sex *k*. While loci contribute additively to the phenotype, alleles within each locus may interact non-additively through dominance.

At locus *ℓ* ∈ {1, 2, … , *L*}, each allele *i* ∈ {1, 2, … , *n*_*ℓ*_} is characterized by an allelic value *x*_*i,ℓ*_, which gives the contribution of that locus when homozygous, such that *z*_*ℓ,k*_ (*x*_*i,ℓ*_, *x*_*i,ℓ*_) = *x*_*i,ℓ*_ in both sexes. For heterozygotes carrying alleles *i* and *j*, the contribution of locus *ℓ* lies between *x*_*i,ℓ*_ and *x*_*j,ℓ*_, with the exact value determined by dominance.

To allow sex-specific dominance to evolve, we use the promoter-affinity framework of Van Dooren (1999), specifically the independent-affinity model of Siljestam et al. (2024). In addition to its allelic value *x*_*i,ℓ*_, each allele carries two sex-specific promoter affinities, *α*_*i,ℓ*, ♀_ and *α*_*i,ℓ*,♂_ , describing its relative recruitment of transcriptional machinery in females and males. Under a fixed total transcriptional output at each locus, the affinities determine the proportional contribution of each allele to the heterozygote phenotype. The contribution of locus *ℓ* to the phenotype of sex *k* is therefore

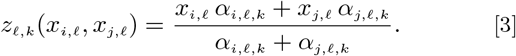

Because promoter affinities cancel in homozygotes, they do not affect the homozygote phenotype and are therefore exposed to selection only at heterozygous loci. Equal promoter affinities within a sex, *α*_*i,ℓ,k*_ = *α*_*j,ℓ,k*_ , correspond to additive allelic effects at that locus. If *α*_*i,ℓ,k*_ ≫ *α*_*j,ℓ,k*_ , allele *i* is nearly completely dominant in sex *k*, whereas if *α*_*i,ℓ,k*_ ≪ *α*_*j,ℓ,k*_ , allele *i* is nearly completely recessive. Dominance is sex-specific whenever the ratio of promoter affinities differs between females and males.

For two alleles ordered such that *x*_*i,ℓ*_ *> x*_*j,ℓ*_, we define their difference in allelic value as Δ*x* = *x*_*i,ℓ*_ − *x*_*j,ℓ*_ *>* 0 and their difference in log-affinity in sex *k* as

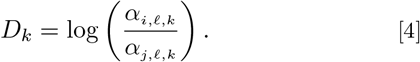

The corresponding dominance weight of allele *i* is 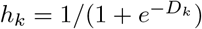, such that *z*_*ℓ,k*_ = *h*_*k*_*x*_*i,ℓ*_ + (1 − *h*_*k*_)*x*_*j,ℓ*_. Thus, *D*_*k*_ *>* 0 corresponds to dominance of the allele with the higher allelic value *x* in sex *k*. We refer to *D*_♂_ *> D*_♀_ as beneficial sex-specific dominance: the higher-valued allele contributes relatively more in males than in females, and equivalently, the lower-valued allele contributes relatively more in females. Complete beneficial dominance reversal is the limiting case in which the lower-valued allele is dominant in females and the higher-valued allele is dominant in males.

This model reduces to the single-locus model of Siljestam et al. (2024) when *L* = 1. For *L >* 1, it allows us to ask whether the feedback between SA selection, sex-specific dominance, and allelic diversification persists when a shared trait is encoded by many independently segregating loci of smaller effect. In particular, we ask whether beneficial sex-specific dominance can maintain allelic polymorphism across a polygenic architecture and how the resulting phenotypic effects become distributed among loci.

### Analysis

#### Analytical expectations

We use evolutionary invasion analysis (Metz et al., 1992; Geritz et al., 1998; Kisdi and Geritz, 1999; Doebeli, 2011) to derive expectations for the initial evolution of the polygenic trait. This approach describes evolution through the invasion success of rare mutant alleles when mutational effects are small and population dynamics occur on a faster timescale than mutation. Although adaptive dynamics was originally developed for traits encoded by a single locus, it has subsequently been extended to diploid and multilocus traits (Geritz et al., 1998; Kisdi and Geritz, 1999; Van Dooren, 1999; Metz and de Kovel, 2013).

When mutations are rare and of small effect, evolution initially proceeds through a sequence of allelic substitutions while the population remains close to monomorphic at the selected loci. A mutant allele arising at one locus either goes extinct or fixates, generating an allelic substitution sequence (Dercole and Rinaldi, 2008; Priklopil and Lehmann, 2020; Champagnat et al., 2006). During this initial, nearly monomorphic phase, a mutation at any focal locus therefore changes the phenotype against an otherwise approximately fixed genetic background. For a focal locus *ℓ*, let

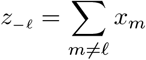

denote the phenotypic contribution of all other loci, such that *z* = *z*_−*ℓ*_ + *x*_*ℓ*_. The background contribution *z*_−*ℓ*_ merely adds a constant offset to the individual’s phenotype for any given value of *x*_*ℓ*_, without changing the strength of selection *σ*^−1^ and *σ*^−1^ or the distance *δ* between the optima. Mutations at the focal locus therefore experience the same local selection as in the corresponding single-locus model (Metz and de Kovel, 2013). Consequently, the single-locus results of Siljestam et al. (2024) for the initial evolutionary dynamics up until the potential emergence of disruptive selection apply locally to mutations arising at any of the *L* loci. We summarize these consequences here and give the formal multilocus argument in Supplementary Material S1.

Gradual evolution initially leads the phenotype toward the singular attractor

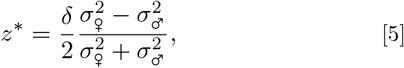

which maximizes geometric mean fitness across the two sexes (Flintham et al., 2024; Siljestam et al., 2024). For a given background contribution *z*_−*ℓ*_, the corresponding singular allelic value at the focal locus is 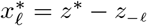. Thus, unlike in the single-locus model, there is no unique combination of allelic values corresponding to the singular phenotype. Rather, any monomorphic configuration satisfying

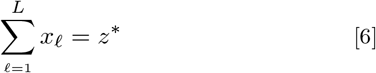

produces the same singular phenotype.

From the single-locus model (Siljestam et al., 2024) follows that, at 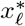(equivalently, when *z* = *z*^∗^), two nearby alleles at a focal locus experience divergent selection when

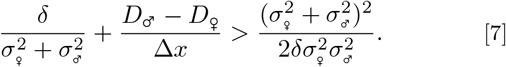

Under symmetric selection strength, *σ*_♀_ = *σ* = *σ*_♂_, this simplifies to

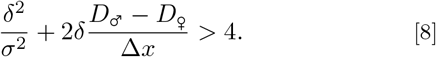

These conditions allow for two immediate observations. First, in the absence of sex-specific dominance, *D*_♀_ = *D*_♂_, they reduce to the evolutionary branching condition derived previously for additive allelic effects (Flintham et al., 2024; Siljestam et al., 2024). Nearby allelic lineages therefore diverge only when selection is sufficiently strong and symmetric. Under symmetric selection, this requires

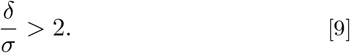

For *δ* = 1, the boundary *σ*^−1^ = 2 corresponds to the locally linear fitness trade-off illustrated in Figure 1, while the focal simulations with *σ*^−1^ = 1 and *σ*^−1^ = 3 lie on either side of this threshold. Second, beneficial sex-specific dominance, *D*_♂_ *> D*_♂_, makes divergence more permissive. Because its contribution scales with (*D*_♂_ − *D*_♀_)*/*Δ*x*, only a small difference in sex-specific dominance is required when the mutant and resident alleles are similar. Heterozygosity can therefore initiate the same positive feedback identified in the single-locus model: beneficial sex-specific dominance can turn stabilising selection toward 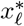 into divergent selection, while the resulting divergence in *x* strengthens selection for further sex-specific dominance (Siljestam et al., 2024). Thus, alleles segregating at a locus can begin to diverge even in parameter regions where selection is stabilising under additivity.

During this initial, nearly monomorphic phase, the same local conditions apply to mutations arising at any of the contributing loci. Which loci first develop diverging polymorphism nevertheless depends on the stochastic order in which mutations arise and establish. Once polymorphism becomes appreciable at several loci simultaneously, mutations occur on heterogeneous genetic backgrounds and the single-locus reduction no longer applies (van Doorn and Dieckmann, 2006).

Under additive allelic effects, such multilocus interactions are known to destabilize symmetric divergence across loci and instead concentrate genetic variation in a small number of loci. In a setting with two phenotypic optima, variation will ultimately become concentrated at a single locus (van Doorn and Dieckmann, 2006; Kopp and Hermisson, 2006). We therefore use individual-based simulations to investigate the subsequent dynamics once substantial multilocus polymorphism and sex-specific dominance have evolved.

We additionally derive analytical expectations for the multilocus endpoint, comparing alternative genetic architectures in Supplementary Material S2. We compare a single-locus biallelic polymorphism (BAP) with multilocus architectures in which all loci are polymorphic with beneficial sex-specific dominance generating sexual dimorphism. We derive the relationship between sexual dimorphism and within-sex phenotypic variance, and obtain an approximate threshold at which a distributed multilocus architecture attains higher mean fitness than the single-locus architecture. These predictions are compared with the evolutionary outcomes of the simulations below.

#### Simulations

We investigate the polygenic evolutionary dynamics of sex-specific dominance and allelic values *x* using individual-based simulations based on Wright–Fisher population dynamics with mutation and selection (Fisher, 1930; Wright, 1931). We use a population size of *N* = 5 *×* 10^4^ and let the trait be encoded by *L* = 30 loci (except for Figure 7, where we consider all numbers from 1 to 30). Simulations additionally include 20 neutral loci that do not affect the phenotype, providing a reference for allelic diversity maintained by mutation and drift alone.

Mutations can affect either allelic values *x*_*i,ℓ*_ or promoter affinities *α*_*i,ℓ,k*_ . Mutational effects on allelic values *x* are drawn from a normal distribution with expected step size 0.03, with per-locus mutation rate *µ*_*x*_ = 5 *×* 10^−6^. Mutations affecting promoter affinities act additively on log(*α*_*i,ℓ,k*_), corresponding to proportional changes in affinity. Mutational effects on female and male log-affinities are drawn jointly from a bivariate normal distribution with expected marginal step size log(2) and correlation *ρ*. Unless stated otherwise, we use *ρ* = 0.5. The per-locus promoter mutation rate is *µ*_*α*_ = 5*×*10^−5^. Promoter affinities are initialized at *α*_*i,ℓ,k*_ = 1 and restricted to *α*_*i,ℓ,k*_ ∈ [10^−2^, 10^2^], with mutational values falling outside these bounds truncated at the corresponding boundary. Simulations are initialized with all selected loci monomorphic and contributing equally to the initial phenotype, *x*_*ℓ*_ = *z*_0_*/L*. We focus first on four simulations that contrast weak and strong symmetric selection, with and without promoter-affinity evolution (Figs. 2–5). Here, time is shown on a log scale and mutations are turned off after 5 *×* 10^5^ generations, indicated by the dashed grey line. The width of each locus-specific trajectory is proportional to allele count. For these simulations, *z*_0_ = −1 and *z*^∗^ = 0. In the parameter sweeps shown in Figures 6 and 7, the initial phenotype is instead placed 0.5 units from the corresponding *z*^∗^.

**Figure 2.**
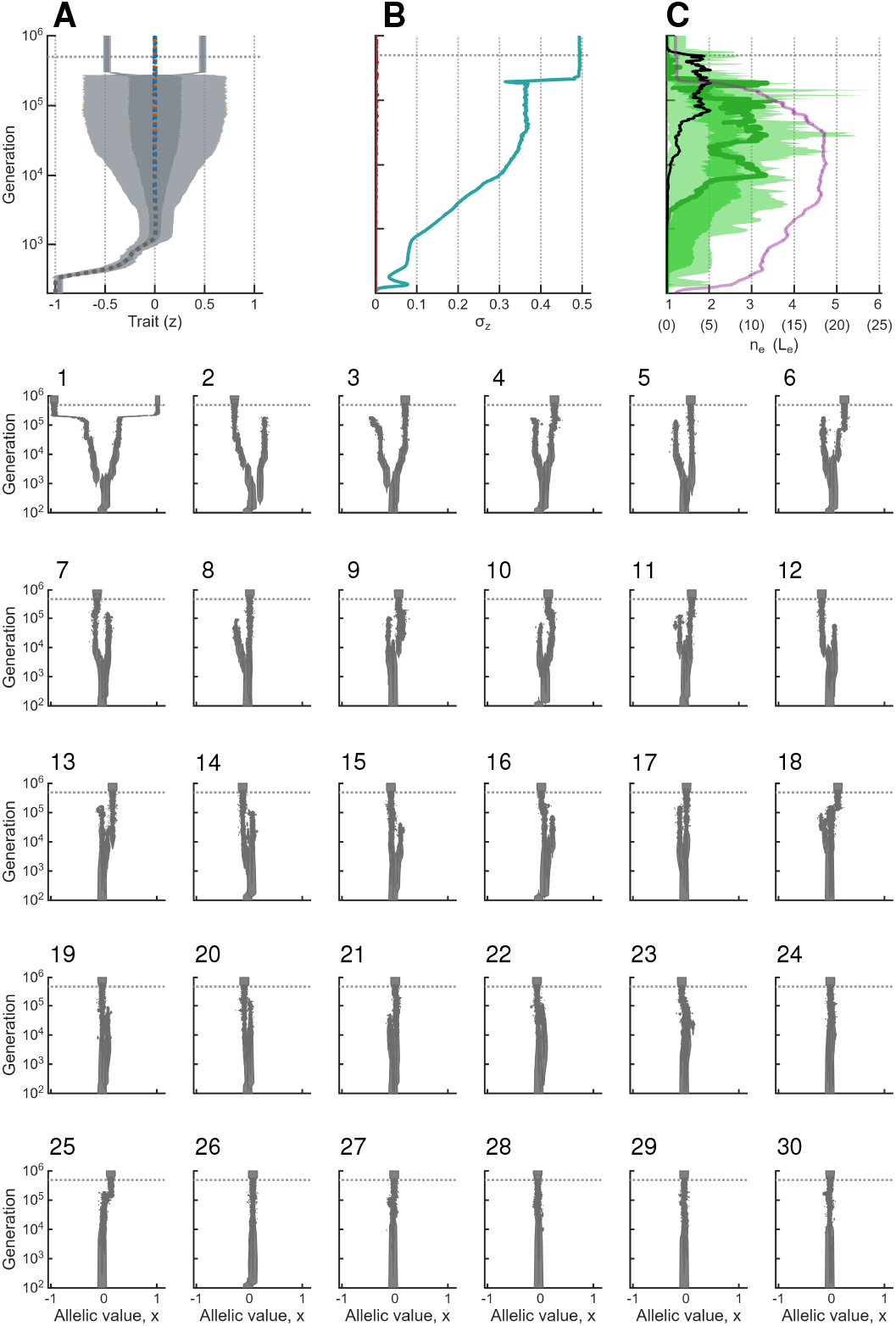
Individual-based simulation under additive allelic effects and strong selection, 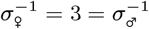. Panels 1–30 show evolutionary trajectories at the 30 selected loci. Panel **A** shows the phenotypic distribution, with darker and lighter shading giving the 50% and 95% contours and dashed lines indicating the mean phenotype. Panel **B** shows the within- and between-sex phenotypic standard deviations, *σ*_*z*,w_ and *σ*_*z*,b_, in teal and red, respectively; the latter is zero in the absence of sexual dimorphism. Panel **C** shows the distribution and weighted mean of the effective number of alleles *n*_*e*_ across selected loci as green shading and a solid green line, respectively, with the mean weighted by *σ*_*z,ℓ*_. The effective number of contributing loci *L*_*e*_ is shown in purple, and the mean *n*_*e*_ across 20 neutral loci in black.

The four focal simulations run for 5 *×* 10^5^ generations with mutation followed by 5 *×* 10^5^ generations without mutational input. Parameter-sweep simulations run for 10^5^ generations with mutation followed by another 10^5^ generations without mutation. The mutation-free phase allows us to distinguish polymorphism maintained by balancing selection from variation maintained only by recurrent mutation. At the end of this phase, we record the number of segregating alleles differing in *x* at each locus.

Furthermore, we calculate the effective number of alleles at each locus as 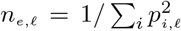, where *p*_*i,ℓ*_ is the frequency of allele *i* at locus *ℓ*. For the parameter sweeps, we additionally calculate the mean number of alleles across loci, weighted by their contribution to phenotypic variation, 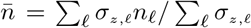, where 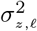 is the variance in the phenotypic contribution of locus *ℓ* across individuals.

We also record the total phenotypic variance of the population, 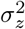, which can be partitioned into within- and between-sex components,

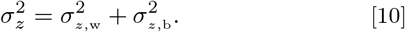

Here, 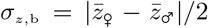 measures the degree of sexual dimorphism, giving the standard deviation between the sex-specific phenotypic means, while *σ*_*z*,w_ gives the standard deviation of phenotypic variation around the sex-specific means.

To quantify the effective number of loci *L*_*e*_ contributing to phenotypic variation, we calculate an inverse-Simpson effective number using the standard-deviation-scale contribution of each locus, *σ*_*z,ℓ*_. Thus,

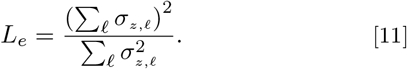

This quantity equals *L* when all loci contribute equally to phenotypic variation and approaches one when phenotypic variation is concentrated at a single locus.

## Results

Our results consist of two parts. First, we show that the single-locus dynamics described in Siljestam et al. (2024) generalize to the initial evolution of the polygenic model: directional evolution approaches the same phenotypic attractor *z*^∗^, while the coevolutionary feedback between sex-specific dominance and allelic divergence can initiate locally at any contributing locus. Under sufficiently strong and symmetric selection, divergence can also initiate through evolutionary branching once *z*^∗^ is reached.

Second, we show that new evolutionary phenomena emerge specifically because the trait is encoded by many loci, generating genetic architectures that differ qualitatively depending on whether sex-specific dominance can evolve. Under additive allelic effects, initially distributed polymorphism ultimately becomes concentrated in a single locus, in agreement with results by van Doorn and Dieckmann (2006) and Kopp and Hermisson (2006). With evolving sex-specific dominance, polymorphism and phenotypic effects instead evolve to become evenly distributed across loci. Increasingly strong selection promotes higher allelic diversity within each locus. We illustrate these dynamics in Figures 2–5, showing evolution across 30 loci under strong and weak symmetric selection, with and without promoter-affinity evolution.

### Initial evolution and onset of diversification

In all four focal simulations, the phenotype initially evolves toward *z*^∗^ = 0 through mutations increasing allelic values at multiple loci (panel **A** in Figs. 2–5). Because successful mutations arise at several loci during this directional phase, segregating variation is already present at multiple loci by the time the population approaches *z*^∗^.

Under strong selection, 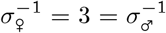, the branching condition (Condition 9) is fulfilled. As the population approaches *z*^∗^, this condition applies locally at each locus: nearby mutant alleles can therefore invade and coexist with resident alleles, with disruptive selection subsequently favouring their divergence. This diversification in *x* does not require the evolution of promoter affinities (Fig. 2), although divergence becomes established asynchronously across loci because mutations arise and spread stochastically. Initial divergence across multiple loci likewise occurs when promoter affinities are allowed to evolve (Fig. 3).

**Figure 3.**
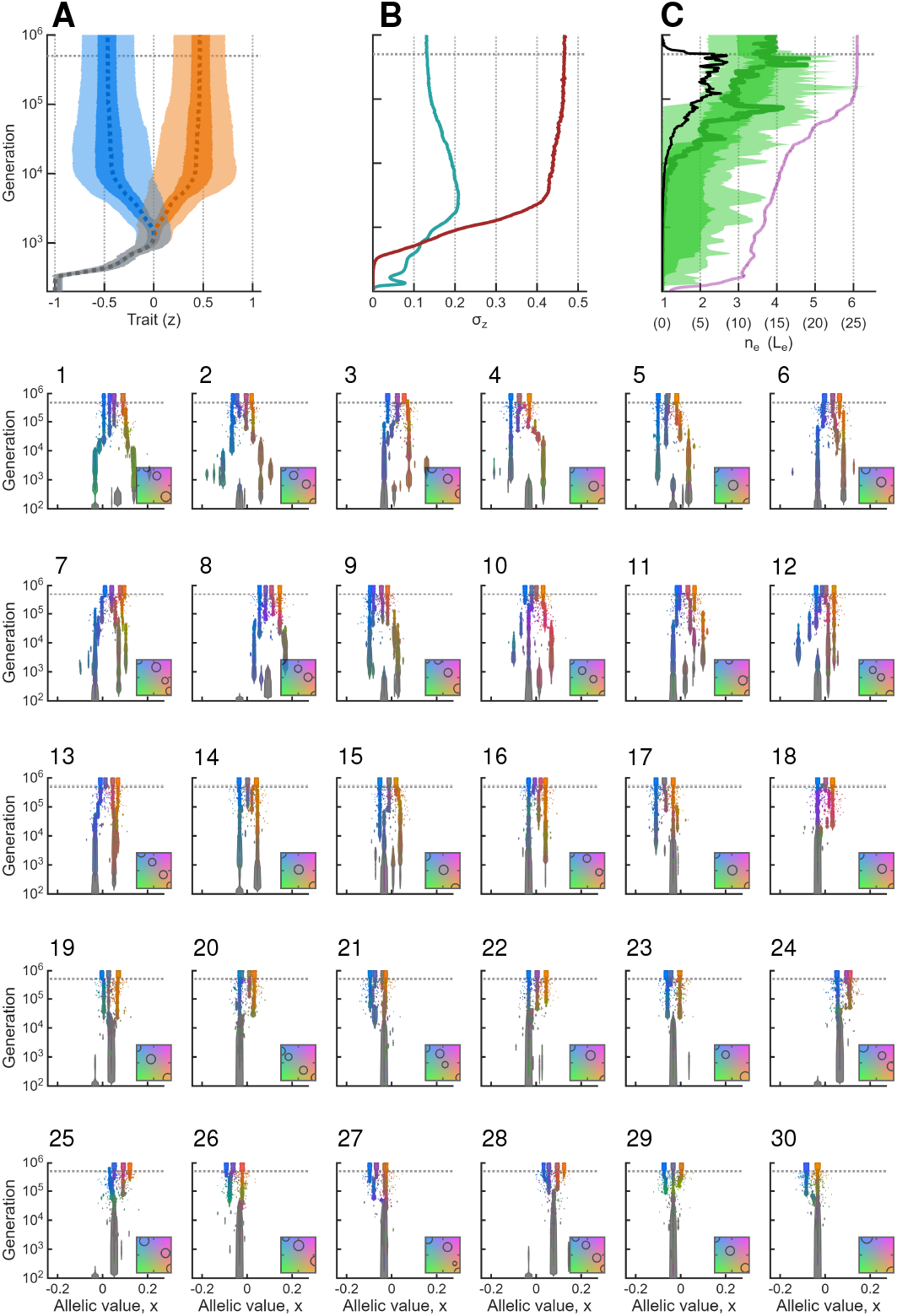
Individual-based simulation under evolving dominance and strong symmetric selection, 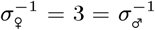. Colours in panels 1–30 indicate the sex-specific promoter affinities *α*_♀_ and *α*_*♂*_ according to the inset colour scale. Panel **A** shows female and male phenotypic distributions in blue and orange, respectively, with grey indicating overlap; dashed lines give the sex-specific means. Panels **B–C** show the same quantities as in Figure 2.

Under weak selection, 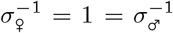, the branching condition 9 is not fulfilled. With fixed promoter affinities, allelic variation therefore converges toward a monomorphic configuration satisfying 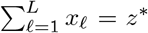 (Fig. 4). Allowing promoter affinities to evolve changes this outcome. Segregating variation provides heterozygotes on which selection on promoter affinities can act, allowing beneficial sex-specific affinity differences to evolve at polymorphic loci (Fig. 5). As predicted by Condition 7, beneficial sex-specific dominance can then change locally convergent selection on allelic values into divergent selection. The resulting positive feedback between sex-specific dominance and divergence in *x* allows polymorphism to become established across multiple loci even under weak selection (Fig. 5; see also Siljestam et al., 2024 for the single-locus mechanism).

**Figure 4.**
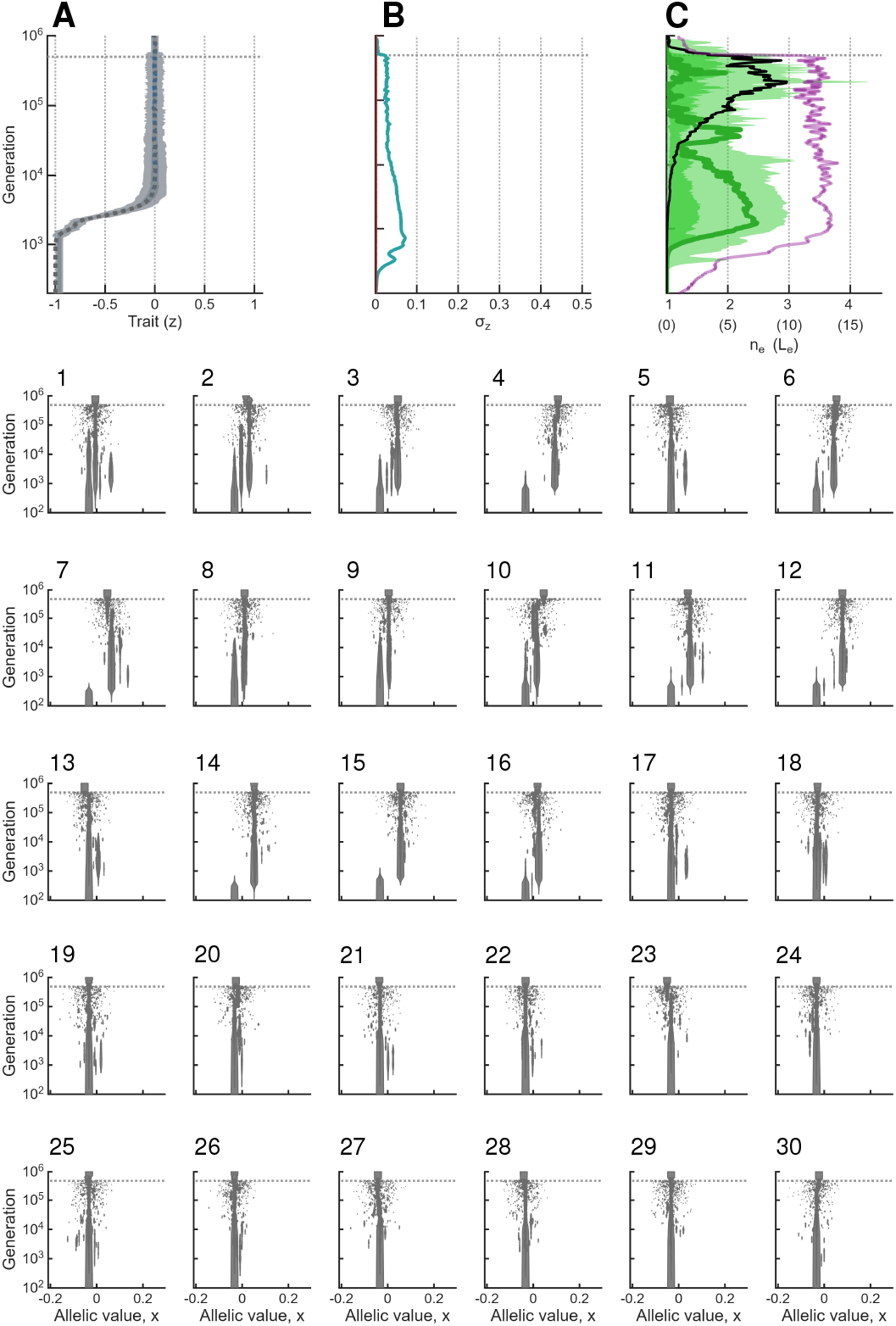
Individual-based simulation under additive allelic effects and weak selection, 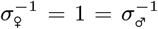. For this selection strength, the branching condition 9 is not fulfilled. See caption of Figure 2 for additional details.

**Figure 5.**
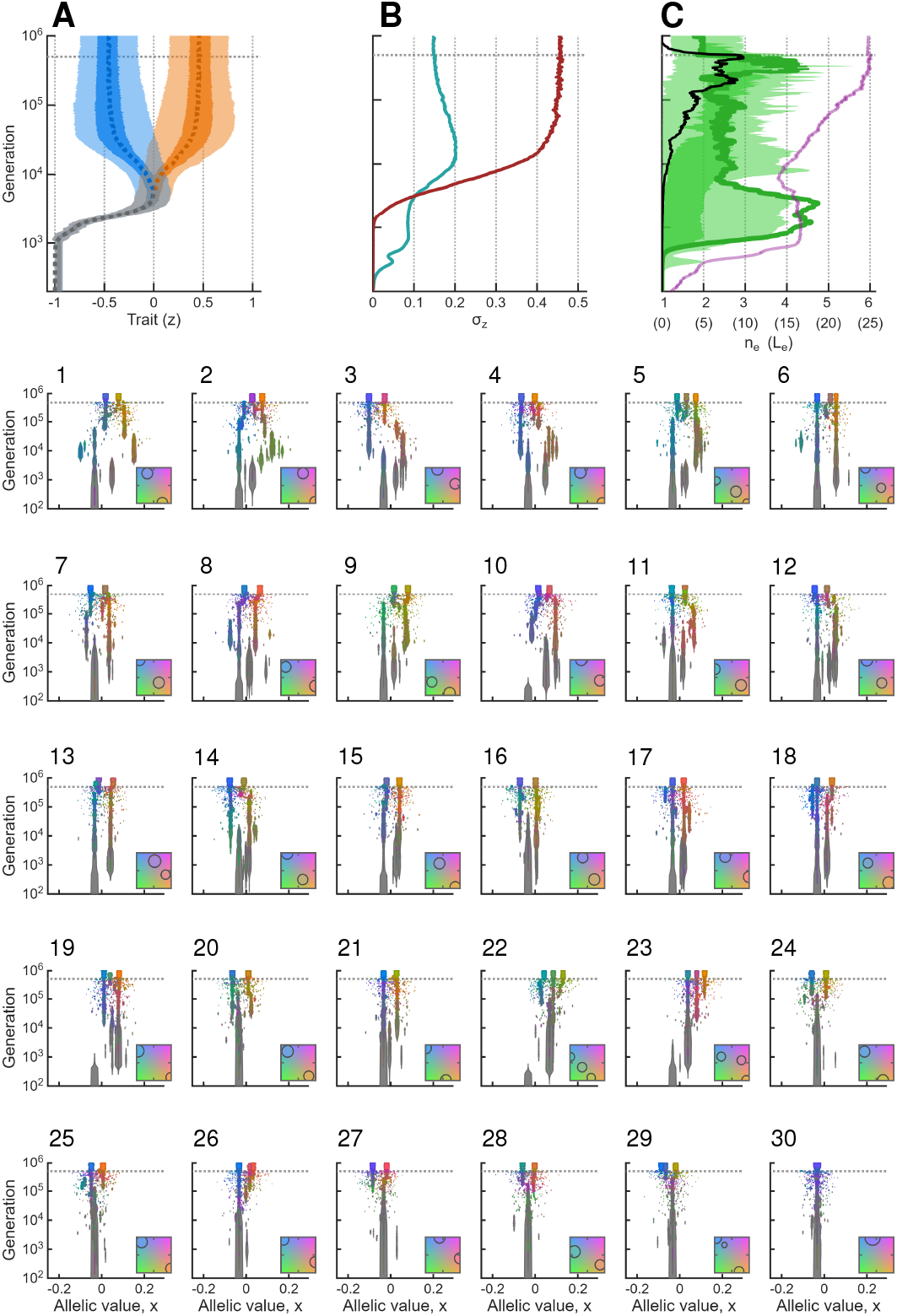
Individual-based simulation under evolving dominance and weak symmetric selection 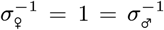. See caption of Figure 3 for additional details.

As beneficial sex-specific dominance evolves across multiple loci, the female and male phenotypic distributions begin to separate (panel **A** in Figs. 3 and 5), accompanied by an increase in *σ*_*z*,b_ (red lines in panels **B**).

### Multilocus evolution after diversification

The single-locus analysis predicts whether individual loci initially experience convergent or divergent selection, but it does not determine the subsequent fate of polymorphism once several loci have diverged. At this stage, selection at any one locus depends on the allelic states and dominance relationships at the remaining loci, and the evolutionary dynamics therefore become genuinely multilocus.

Under strong selection (Figs. 2–3), polymorphism already arises at several loci as the population approaches *z*^∗^. The key contrast between fixed and evolving promoter affinities therefore concerns the subsequent fate of this initially distributed variation.

Without promoter evolution, divergence at multiple loci initially broadens the phenotypic distribution around *z*^∗^ while the distribution remains unimodal (Fig. 2**A**). Although allelic divergence generates some genotypes that approach the female and male optima, recombination continually produces mostly intermediate phenotypes, which are poorly adapted. As allelic divergence proceeds, the initially distributed polymorphism becomes unstable: differentiation continues to increase at the most divergent loci, while polymorphisms at other loci collapse one after another. Ultimately, all phenotypic differentiation becomes concentrated at a single polymorphic locus (panels 1–30 in Fig. 2). This reproduces the characteristic concentration of polymorphism found by van Doorn and Dieckmann (2006) in their closely related multilocus model of adaptation to two habitats and is consistent with the more general tendency for disruptive selection to concentrate phenotypic effects in a small number of loci (Kopp and Hermisson, 2006).

As polymorphism collapses, loci become fixed for either female- and male-beneficial allelic values, keeping their combined contribution close to *z*^∗^. This corresponds to the “equity effect” described by Arnqvist et al. (2014), whereby fixation benefiting one sex at one locus shifts selection at other loci toward alleles benefiting the opposite sex.

The remaining polymorphic locus evolves such that one homozygote and the heterozygote approximately match the two sex-specific optima, while the second homozygote lies beyond one optimum. Reproduction then occurs almost entirely through the locally adapted homozygote in one sex and the heterozygote in the other. Their crosses produce the two high-fitness genotypes in approximately equal proportions, while the maladapted homozygote is produced at negligible frequency and therefore does not appear within the 95% contour of the phenotypic distribution in Figure 2**A**. This pattern also occurs in the corresponding single-locus model (e.g., Fig. 4**iii** in Siljestam et al., 2024), and is mirrored in a single-locus model of habitat specialization (Kisdi and Geritz, 1999).

With evolving promoter affinities, the fate of polymorphism is qualitatively different. We illustrate this in Figure 3 for strong and Figure 5 for weak selection. Beneficial sex-specific dominance evolves at polymorphic loci, generating distinct sex-specific genotype-to-phenotype maps at heterozygous loci. The female and male phenotypic distributions consequently separate, such that the pooled phenotypic distribution becomes bimodal, with the mean of each sex approaching its respective optimum (panel **A** in Figs. 3 and 5). This increasing sexual dimorphism is reflected by the increase in *σ*_*z*,b_ (panel **B**).

Once the female and male phenotypic means approach their respective optima, phenotypic effects are redistributed across loci: initially strongly differentiated loci decrease in effect, while loci with little or no differentiation develop diverging polymorphisms, until all 30 loci become polymorphic (panels 1–30). Consequently, *L*_*e*_ increases toward the actual number of selected loci (panel **C**), and the locus-specific contributions to phenotypic variation become smaller and more even.

This redistribution causes a decline in within-sex phenotypic variation, *σ*_*z*,w_ (teal line in panel **B**), resulting in increasingly narrow female and male phenotypic distributions around their respective optima (panel **A**). At the same time, sexual dimorphism, measured by *σ*_*z*,b_, continues to increase. Thus, distributing the genetic basis of sexual dimorphism across more loci reduces the phenotypic variation among individuals within each sex. The analytical model gives the same result: for an idealized distributed architecture generating a fixed degree of sexual dimorphism, within-sex phenotypic variance decreases as 1*/L* (Supplementary Material S2).

Allelic diversity also increases within the already distributed set of polymorphic loci. More loci evolve polyallelic polymorphisms (PAPs) accompanied by sex-specific dominance hierarchies, visible as multiple differently coloured allelic lineages within loci and by the distribution of *n*_*e*_ shifting above two (panel **C**). Under stronger selection, PAPs occur at more loci (compare Figs. 3 and 5). Notably, this differs from the single-locus model, in which PAP was favoured primarily under weak selection (Siljestam et al., 2024). In the multilocus model, the range of selection strengths over which a distributed architecture is favoured increases with the number of contributing loci, consistent with the analytical comparison in Supplementary Material S2 (Eq. S45), and stronger selection instead promotes increasing allelic diversity within the already polymorphic loci. This produces an additional reduction in within-sex variance and consequently still narrower phenotypic distributions around the sex-specific optima (panels **A–B**). The analytical *n*-allele PAP model in Supplementary Material S2 shows the same effect: for a fixed degree of sexual dimorphism, within-sex variance decreases as the number of alleles in the dominance hierarchy increases (Eq. S27).

#### Dependence on selection strength and number of loci

Figure 6 extends these results across combinations of female and male selection strength for a trait encoded by *L* = 30 loci. Without promoter-affinity evolution (i.e., under additivity, left column in Fig. 6), polymorphism is largely restricted to the region in which the evolutionary branching condition is fulfilled (Condition 7 with *D*_♀_ = *D*_♂_), that is, under strong and symmetric selection. Long-term phenotypic variation then becomes concentrated in a small number of loci, as in Figure 2. The hatched white line in column **A** in Figure 7 shows that this outcome is largely insensitive to the number of loci and only depends on whether the condition for evolutionary branching is fulfilled.

Allowing promoter affinities to evolve produces persistent polymorphism across the full parameter space. Phenotypic effects are distributed across loci, with *L*_*e*_ approaching *L* over most of the parameter space (Fig. 6**B**). The number of alleles within these loci responds differently: 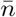 remains close to two under weak selection but increases progressively as selection becomes stronger in either sex. Values of 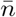 above two indicate polyallelic polymorphism among the loci contributing to phenotypic variation. Thus, for *L* = 30 and over the range of selection strengths shown here, stronger selection primarily increases allelic diversity within an already distributed polymorphism.

**Figure 6.**
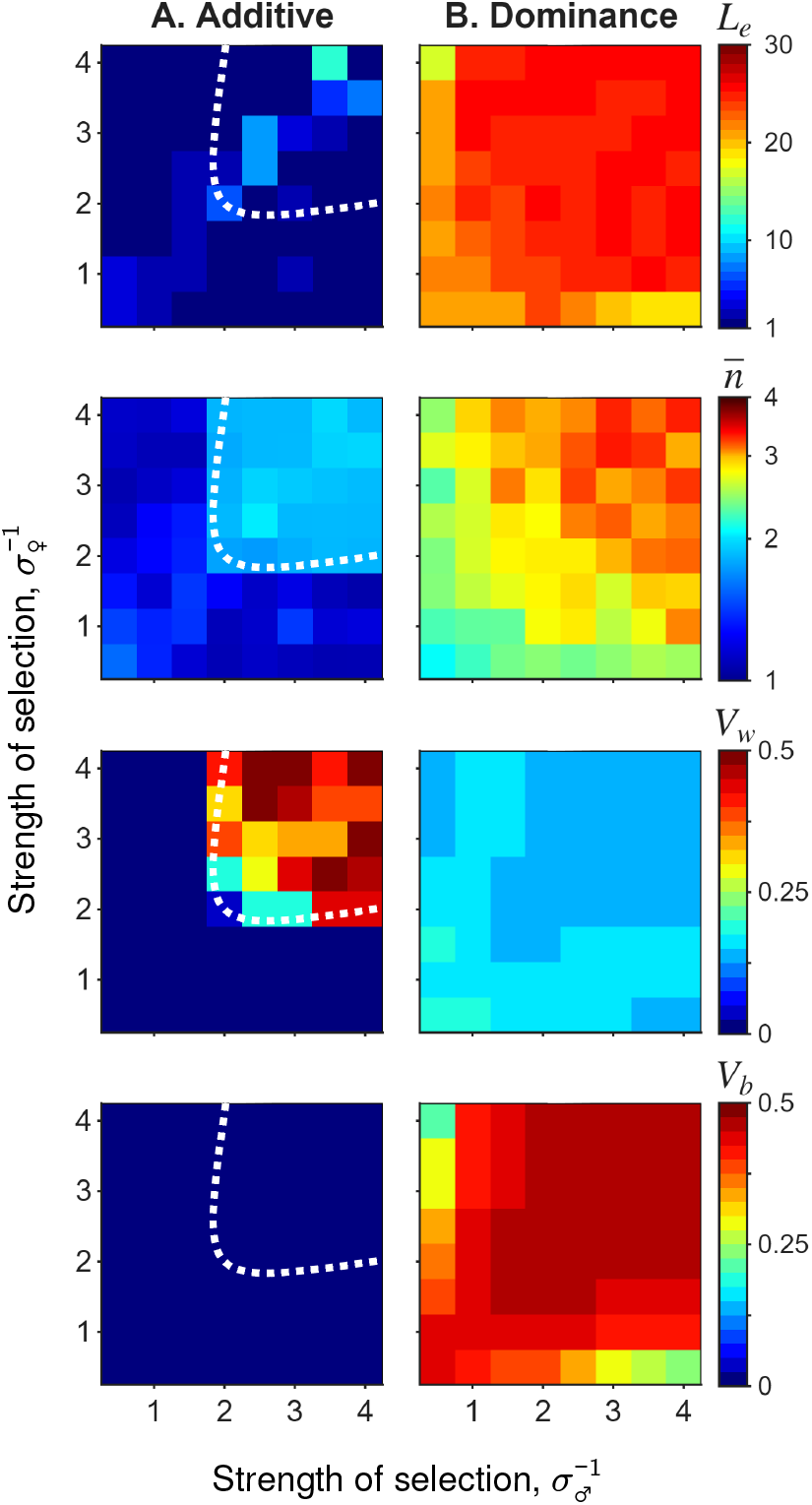
Evolutionary outcomes as a function of the strength of selection in the two sexes (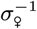 and 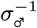) derived from individual-based simulations with *L* = 30 selected loci. Columns show results for **A**, no affinity evolution, and **B**, evolving promoter affinities. Each pixel represents the outcome of a single simulation. The four rows show the effective number of loci (*L*_*e*_) contributing to phenotypic variation; the mean number of alleles 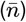 across loci, weighted by the locus-specific standard deviation *σ*_*z,ℓ*_; within-sex phenotypic variance 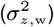; and between-sex phenotypic variance 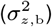. The dotted white lines in column **A** indicate the analytical threshold for evolutionary branching under additive allelic effects (Flintham et al., 2024). Simulations are initialized with *z*_0_ = *z*^∗^ ∑0.5, ensuring comparable initial directional selection across parameter combinations.

This pattern eventually breaks down under substantially stronger selection, and the strength required for this transition depends strongly on the number of loci available to encode the trait. As the number of loci decreases or selection becomes very strong, the within-sex segregation variance generated by the distributed polymorphism becomes increasingly detrimental relative to the single-locus alternative, causing polymorphism to collapse toward a single major locus. This is explored in Figure 7, where the polymorphism collapse occurs at approximately *σ*^∑1^ *>* 2.5 for *L* = 2 and *σ*^∑1^ *>* 4 for *L* = 10, whereas for *L* = 30 collapse occurs only around *σ*^∑1^ ≳ 6.5. Thus, the selection-strength sweep in Figure 6, which extends only to 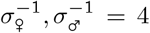, does not reach the single-locus-collapse regime for *L* = 30.

**Figure 7.**
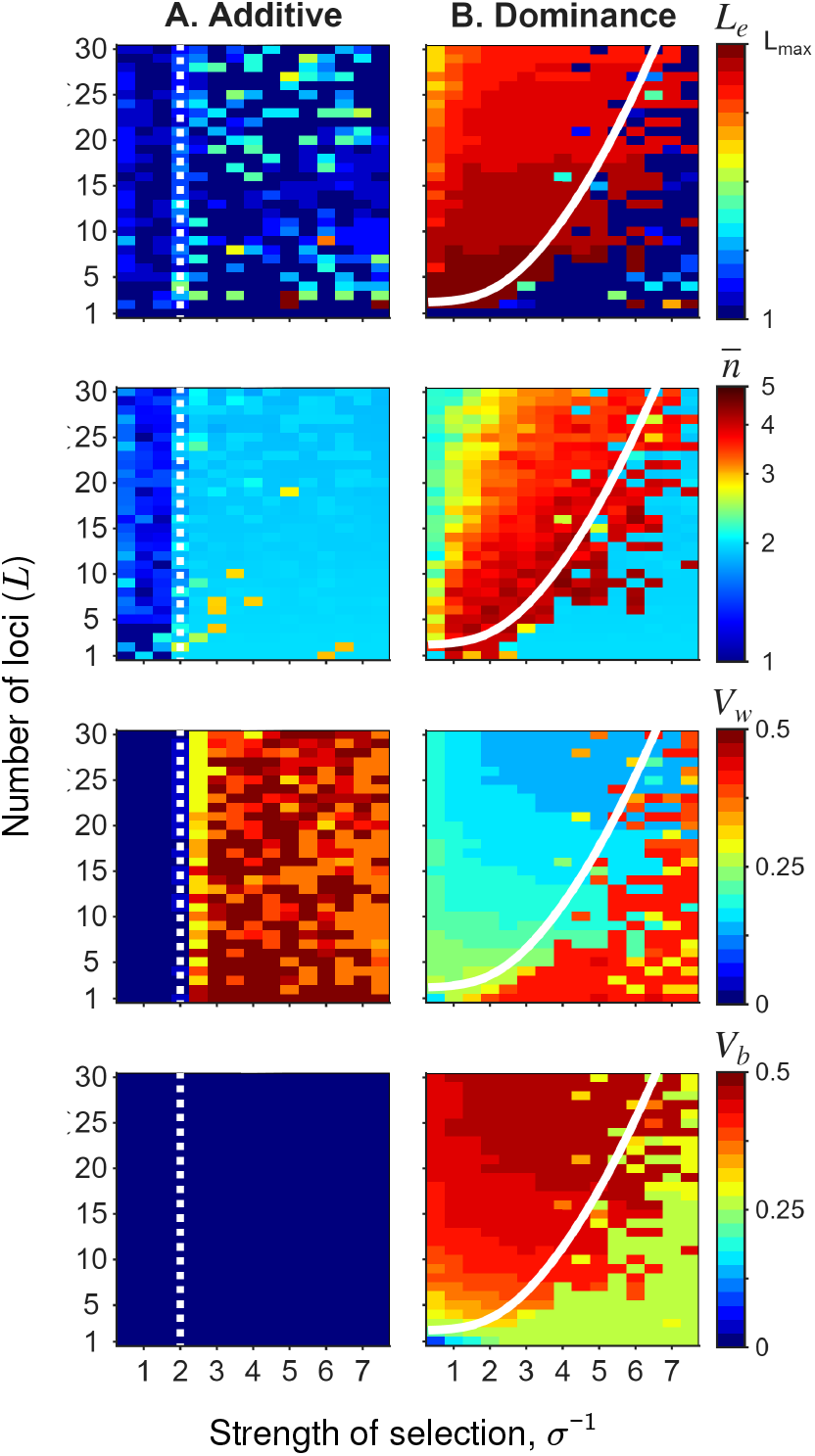
Evolutionary outcomes as a function of the number of selected loci *L* and symmetric selection strength 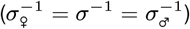, derived from individual-based simulations. Columns show results for **A**, no affinity evolution, and **B**, evolving promoter affinities. Rows show the same quantities as in Figure 6. The solid lines in column **B** give the threshold *L*_crit,4_ at which an idealized multilocus PAP of four alleles is predicted to attain higher mean fitness than the single-locus BAP (Eq. S48). This threshold closely tracks the transition between simulations in which polymorphism collapses toward a single locus and those in which polymorphism and phenotypic effects remain distributed across loci. The dotted white lines in column **A** indicate the threshold for evolutionary branching (Eq. 9). Simulations are initialized with *z*_0_ = ∑ 0.5, displaced by 0.5 from the singular phenotype *z*^∗^ = 0.

The Supplementary Material gives an approximate threshold for this transition by comparing the mean fitness of an idealized four-allele multilocus PAP with that of a single-locus BAP. The resulting *L*_crit,4_ (solid line in column **B** in Fig. 7) closely tracks the transition in the simulations: for parameter combinations with *L* above this threshold, polymorphism is distributed across loci, whereas below the threshold it collapses toward a single locus. Across the two parameter sweeps shown in Figures 6 and 7, increasing selection under evolving dominance therefore has two distinct effects before collapse: *L*_*e*_ remains high while 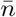 increases, indicating increasing allelic diversity within an already distributed polymorphism. Only at substantially stronger selection does *L*_*e*_ fall sharply as phenotypic variation becomes concentrated at a single locus.

## Discussion

We show that allowing sex-specific dominance to evolve fundamentally changes the genetic architecture favoured by sexually antagonistic (SA) selection acting on a polygenic quantitative trait. Rather than concentrating polymorphism into a single major locus, selection can favour polymorphism distributed across many loci, generating sexual dimorphism while reducing segregation load. Additional alleles within loci reduce this load further, whereas collapse toward a single major locus occurs only under sufficiently strong selection.

Why does a distributed architecture reduce segregation load? A given difference between female and male mean phenotypes can be generated either by large effects at a few loci or by smaller effects distributed across many loci. The latter architecture generates substantially less within-sex phenotypic variance. In our analytical approximation, within-sex variance declines approximately as 1*/L*, while the standard deviation declines as 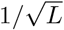. Polyallelic polymorphism and the associated dominance hierarchies reduce this variance further. Female and male phenotypic distributions can therefore become progressively narrower around their respective optima as both the number of contributing loci and the number of alleles within loci increase, and thus allows the genetic conflict between sexes to be progressively resolved.

This result contrasts sharply with multilocus models with additive allelic effects. Under corresponding conditions, we reproduce the outcome found by van Doorn and Dieckmann (2006) and Kopp and Hermisson (2006), who studied polygenic traits under diversifying selection generated by ecological heterogeneity and found that polymorphism initially distributed across loci becomes unstable and genetic variation is eventually concentrated at a single locus. In these scenarios, intermediate phenotypes have lower fitness than more extreme ones and selection favours phenotypic differentiation (Rueffler et al., 2006). Concentrating genetic variation at a single locus reduces the production of intermediate phenotypes by recombination. Our model shows that this outcome is not a general consequence of diversifying selection itself. When genotype-to-phenotype relationships can evolve context-specific dominance, phenotypic dimorphism emerges, and distributing variation across many loci can instead reduce the phenotypic variance within each context, and thus segregation load.

In contrast to our model, van Doorn and Dieckmann (2006) and Kopp and Hermisson (2006) do not allow for the evolution of context-specific dominance. Empirical studies have, however, documented environment-dependent dominance reversal across both temperature and salinity gradients (Chen et al., 2015; Posavi et al., 2014; Fukutomi et al., 2026). Including the evolution of context-specific dominance in corresponding models of ecological diversification would therefore be warranted. Since our model with selection in two sexes is closely analogous to selection across two environments, our results suggest that context-specific dominance can allow disruptive selection to maintain allelic variation across many loci rather than concentrating it into a single major locus.

Our results differ fundamentally from those of Flintham et al. (2026), who analysed a closely related polygenic SA model but found that polymorphism is not maintained over long evolutionary timescales. The difference follows directly from the genotype-to-phenotype map that is allowed to evolve. In their model, the same allele can accumulate different phenotypic effects in females and males. Consequently, selection can produce a homozygous genotype that simultaneously expresses a female phenotype near the female optimum and a male phenotype near the male optimum. Once such a genotype has evolved to match both optima, the sexually antagonistic conflict is removed without any requirement for polymorphism, and alleles maintained through sex-specific dominance are consequently displaced.

Our model excludes this route by assuming that the underlying allelic effect *x* is shared between females and males: a homozygous locus therefore contributes the same value to the phenotype in both sexes. Sex specificity can instead evolve only through the relative expression of alternative alleles in heterozygotes. Under this constraint, polymorphism cannot be bypassed by the evolution of sex-specific allelic effects. Rather, polymorphism itself becomes part of the solution to the conflict. Beneficial sex-specific dominance allows alternative alleles to contribute differently in females and males, and when this mechanism evolves across many loci, selection favours a distributed architecture in which polymorphism is maintained throughout the genetic basis of the trait.

A similar distinction applies to the amplification model of Flintham (2025). There, sex-specific expression can evolve from full expression to complete silencing, and the genetic architectures considered contain sufficient redundancy for the two sexes to use different subsets of the loci underlying the trait. Even at the lowest redundancy considered, males and females can evolve effectively private genomes, with half of the loci contributing only in males and the other half only in females, such that a genotype homozygous at every locus can produce both sex-specific optima. As in the model of Flintham et al. (2026), conflict can thus be eliminated without maintaining allelic polymorphism. Our model addresses the complementary case in which neither sex-specific allelic effects nor sex-limited use of redundant loci can remove the shared genetic constraint.

Our results also differ from the dominance-modifier model of Flintham (2025), in which each locus is restricted to two alleles of fixed phenotypic effect and the evolution of sex-specific dominance becomes increasingly constrained under weak selection as the number of loci increases. In our continuum-of-alleles model, ongoing evolution of allelic values allows the positive feedback between allelic divergence and sex-specific dominance to operate at individual loci, permitting polymorphism and dominance to coevolve even under weak selection.

In summary, allowing for the evolution of sex-specific dominance reverses the usual expectation that diversifying selection should concentrate genetic polymorphism into a small number of major loci. Over a broad range of conditions, polymorphism instead becomes distributed across the polygenic architecture, with stronger selection promoting additional allelic diversity and dominance hierarchies within loci. This architecture allows female and male phenotypes to approach their respective optima while progressively reducing within-sex phenotypic variance. Only when selection becomes strong relative to the number of contributing loci does the segregation load favour a single-locus architecture.

## S1. Initial multilocus adaptive dynamics

Here, we give the formal argument underlying the use of the single-locus invasion conditions in the initial phase of the multilocus model.

Consider a resident population that is monomorphic at all *L* loci for alleles with allelic value *x*_*ℓ*_ at locus *ℓ*. Because promoter affinities do not affect homozygous genotypes, the resident phenotype is

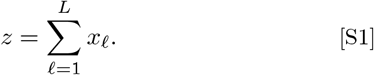

Under the assumption of rare mutations of small phenotypic effect, evolution initially proceeds toward the singular phenotype

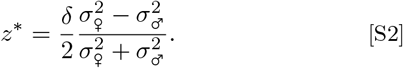

Unlike in the single-locus model, the singular phenotype does not correspond to a unique point in the *L*-dimensional space of allelic values. Instead, all monomorphic configurations satisfying

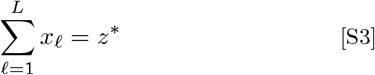

generate the same phenotype and therefore form an (*L* ∑ 1)-dimensional manifold in allele space.

To see why the single-locus invasion condition applies locally at each locus, consider a focal locus *ℓ* and define the contribution of all remaining loci as

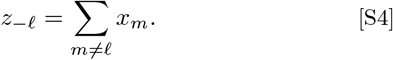

The phenotype can then be written as

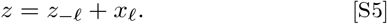

For a mutation arising at locus *ℓ*, the background contribution *z*_∑*ℓ*_ is fixed during the initial invasion. The effective female and male optima for the contribution of the focal locus are therefore 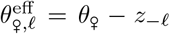 and 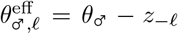.Their separation remains

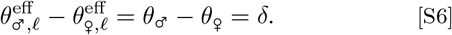

Thus, the background contribution *z*_∑*ℓ*_ only adds a constant offset to the phenotype associated with a given value of *x*_*ℓ*_. It does not change the widths of the fitness functions, *σ*_*♀*_ and *σ*_*♂*_ , or the distance *δ* between the female and male optima. The invasion analysis for a mutation at any focal locus is therefore equivalent to that of the single-locus model after a shift of coordinates. In particular, for a given background *z*_∑*ℓ*_, the singular allelic value of the focal locus is

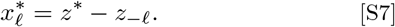

When 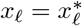, the phenotype satisfies *z* = *z*^∗^.

Consequently, when the population lies on the singular manifold described by Equation S3, then the local conditions for invasion, evolutionary branching, and dominance-mediated divergence derived in Siljestam et al. (2024) apply to mutations arising at each focal locus, as long as standing polymorphism at other loci is sufficiently small. In particular, two nearby alleles at locus *ℓ* experience divergent selection if

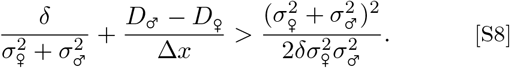

Without sex-specific dominance, this reduces to the additive evolutionary branching condition

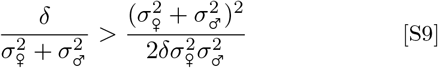

presented by Flintham et al. (2024) and Siljestam et al. (2024). Under symmetric selection, *σ*_♀_ = *σ* = *σ*_♂_ , these conditions reduce to

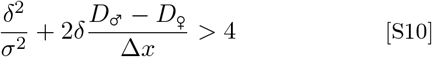

and

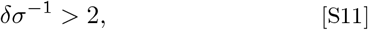

respectively.

This single-locus–multilocus equivalence applies to the initial, nearly monomorphic phase. Once substantial polymorphism is simultaneously maintained at several loci, mutations at a focal locus occur on heterogeneous genetic backgrounds and their invasion success depends on the variation at the other loci. The single-locus reduction then no longer applies, motivating the individual-based simulations used in the main text.

## S2. Single-locus and multilocus limits

We compare a single-locus biallelic polymorphism (BAP) with distributed multilocus architectures containing *n* ≥ 2 alleles per locus. For *n* = 2, the latter is a multilocus BAP, whereas *n >* 2 gives a multilocus polyallelic polymorphism (PAP). We consider idealized symmetric architectures under symmetric selection, *σ*_♀_ = *σ* = *σ*_♂_ , in which alleles occur at equal frequency, loci segregate independently, allelic effects within each locus are symmetric around zero, and complete beneficial sex-specific dominance hierarchies cause females to express the allele with the smaller effect and males the allele with the larger effect at every heterozygous locus. We do not initially assume that allelic effects are equally spaced; this spacing emerges below as the configuration minimizing within-sex phenotypic variance.

The purpose of the following calculation is to determine how the segregation variance associated with a given degree of sexual dimorphism depends on the number of contributing loci and on the number of alleles maintained within each locus. We first derive the minimum within-sex phenotypic variance for such architectures and then use this result to compare their mean fitness with that of a single-locus BAP.

First, consider a single biallelic locus with alleles coding for the female and male optima, ∑*δ/*2 and *δ/*2, respectively, segregating at equal frequency. Under complete beneficial dominance reversal, heterozygotes express the female optimum in females and the male optimum in males. Thus, in each sex, the locally adapted homozygote and the heterozygote have fitness 1, while the maladapted homozygote has fitness

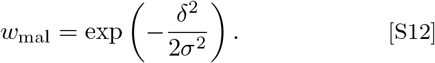

Mean fitness in each sex is therefore

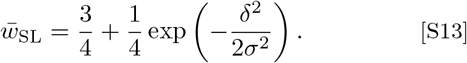

Thus, 3*/*4 of individuals express their sex-specific optimum, whereas 1*/*4 express the opposite maladapted optimum.

We next consider a distributed multilocus architecture with *n* equally frequent alleles at each of *L* loci. At each locus, we order their allelic effects as

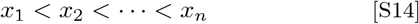

with *x*_*n*+1_∑_*i*_ = ∑*x*_*i*_. Under complete beneficial sex-specific dominance, a male expresses the larger of the two allelic effects carried at a locus, whereas a female expresses the smaller. The probability that allele *i* is expressed in a male is the probability that both sampled alleles have rank at most *i*, minus the probability that both have rank at most *i* ∑ 1. Hence,

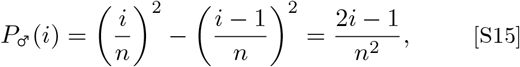

while the corresponding probability in a female is

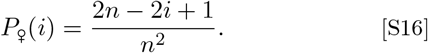

These expressions include homozygotes and therefore apply to both even and odd *n*.

For convenience, define

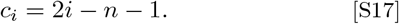

Because the allelic effects are symmetric, ∑_*i*_*x*_*i*_ = 0. The expected contribution of locus *ℓ* to the male phenotype can therefore be written as

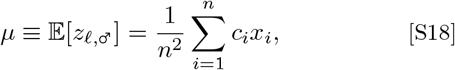

while 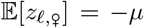. The second moment is the same in both sexes and, by symmetry, simplifies to

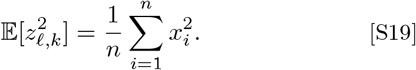

Hence, the within-sex phenotypic variance generated by a single locus is

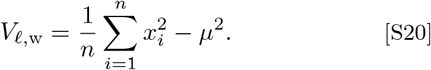

For a fixed difference between the female and male mean phenotypes, *µ* is fixed. Minimizing the within-sex variance is therefore equivalent to minimizing ∑_*i*_*x*^2^ subject to a fixed value of ∑__*i*__ *c*_*i*_*x*_*i*_. From the Cauchy–Schwarz inequality,

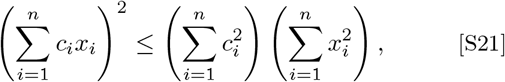

with equality when *x*_*i*_ is proportional to *c*_*i*_. The variance-minimizing allelic effects therefore satisfy

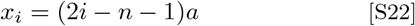

for some *a >* 0. Thus, the optimal allelic effects are equally spaced,

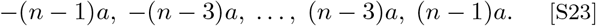

For odd *n*, this sequence includes an allele with effect zero.

Using

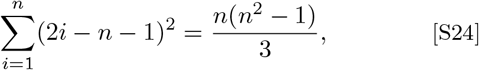

the minimum within-sex variance at one locus can be expressed directly in terms of its sex-specific mean as

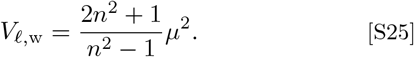

For the symmetric distributed architecture, the difference between the population-wide male and female mean phenotypes is

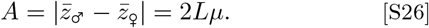

Because loci segregate independently, their variances add, and the total within-sex phenotypic variance is therefore

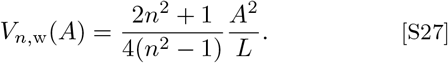

Writing

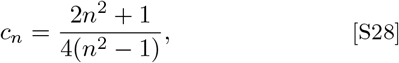

this becomes *V*_*n*,w_(*A*) = *c*_*n*_*A*^2^*/L*.

The multilocus BAP is recovered as the special case *n* = 2, for which *c*_2_ = 3*/*4 and

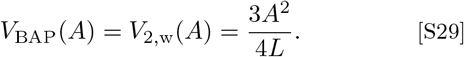

For *n* = 4, *c*_4_ = 11*/*20, giving

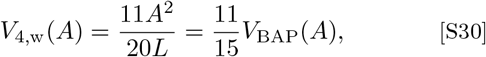

recovering the four-allele result. More generally,

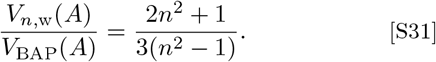

This ratio decreases monotonically with *n* and approaches 2*/*3 as *n*→ ∞ . Thus, increasing allelic diversity progressively reduces the within-sex segregation variance required to generate a given degree of sexual dimorphism, although with diminishing returns.

We next consider how this within-sex variance affects the fitness-maximizing degree of sexual dimorphism. Let

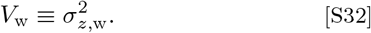

Let the difference between the male and female mean phenotypes be

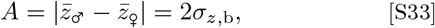

such that

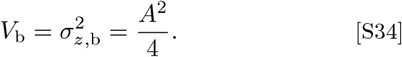

For the symmetric architectures considered here, 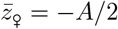 and 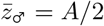.

For large *L*, the phenotype distribution within each sex is approximately Gaussian. If 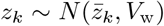 and fitness is *w*_*k*_(*z*_*k*_) = exp[∑(*z*_*k*_ ∑ *θ*_*k*_)^2^*/*(2*σ*^2^)], mean fitness is

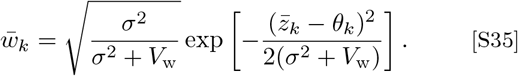

For the multilocus *n*-allele architectures above, the within-sex variance itself depends on the amount of sexual dimorphism according to

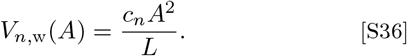

Equation S35 therefore becomes

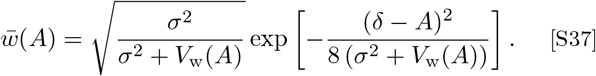

If *V*_w_ were independent of *A*, fitness would be maximized at *A* = *δ*. In a multilocus architecture, however, increasing sexual dimorphism also increases the within-sex phenotypic variance.

Because *V*_*n*,w_(*A*) ∝ *A*^2^ for every *n*, the fitness-maximizing divergence has the same relationship to the within-sex variance for all of these architectures. For the interior optimum relevant here, differentiating Equation S37, using 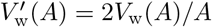, gives

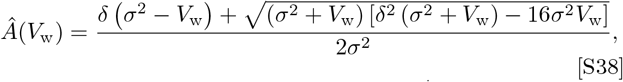

where we take the root approaching *Â* = *δ* as → *V*_w_ 0. Since *V*_b_ = *A*^2^*/*4, the corresponding fitness-maximizing between-sex variance is

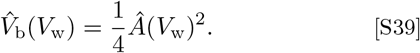

The number of alleles *n* affects how much within-sex variance is generated for a given degree of sexual dimorphism *A*, through the coefficient *c*_*n*_. However, once the realized within-sex variance *V*_w_ is specified, the fitness-maximizing value of *A* depends only on *V*_w_, *δ*, and *σ*, and therefore does not depend explicitly on *n* (Eq. S38).

For small *V*_w_,

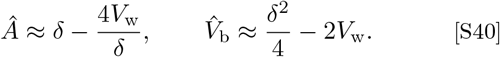

Thus, as within-sex variance decreases, the fitness-maximizing sexual dimorphism approaches the separation between the sex-specific optima.

When the sex-specific means coincide exactly with their optima, *A* = *δ*, Equation S37 reduces to

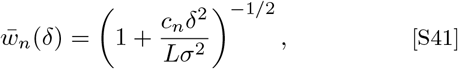

with load

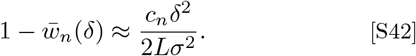

Thus, the segregation load decreases as 1*/L* for any fixed number of alleles *n*. Increasing *n* further reduces this load through the decrease in *c*_*n*_, from *c*_2_ = 3*/*4 for a BAP toward *c*_*n*_ = 1*/*2 as *n* → ∞.

Finally, equating multilocus and single-locus mean fitness gives the maximum within-sex phenotypic variance for which a multilocus architecture centred at the sex-specific optima has at least the same mean fitness as the single-locus BAP:

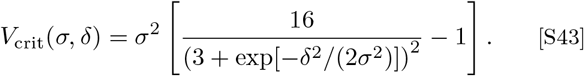

A multilocus *n*-allele architecture therefore has lower load than the single-locus BAP when

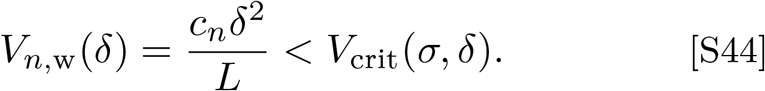

This gives the critical number of loci

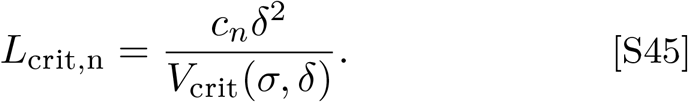

For the multilocus BAP, *n* = 2, this becomes

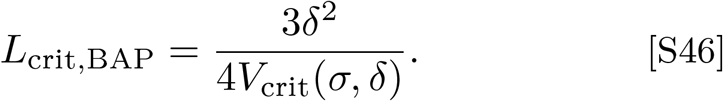

Relative to this BAP threshold,

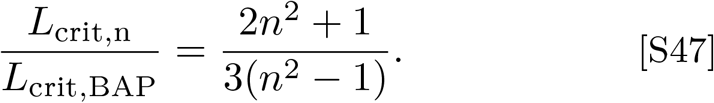

For the four-allele PAP,

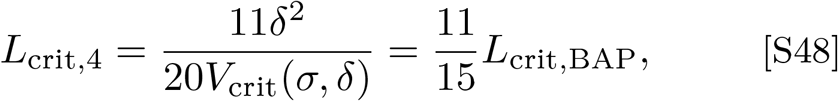

while

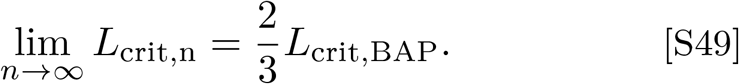

The respective multilocus architecture has lower load when *L > L*_crit,n_. Under weak selection, Equation S45 reduces to

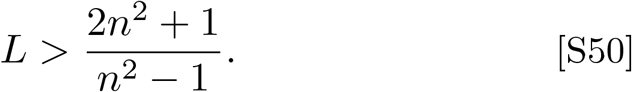

This gives *L >* 3 for the multilocus BAP, *L >* 11*/*5 for the four-allele PAP, and approaches *L >* 2 as *n* → ∞. These thresholds assume *A* = *δ*; allowing *A* to attain its fitness- maximizing value *Â < δ* slightly lowers the thresholds.

